# VirB4- and VirD4-like ATPases, components of a putative type 4C secretion system in *Clostridioides difficile*

**DOI:** 10.1101/2021.07.12.452133

**Authors:** Julya Sorokina, Irina Sokolova, Ivan Rybolovlev, Natalya Shevlyagina, Vasiliy Troitskiy, Vladimir Zhukhovitsky, Yury Belyi

## Abstract

The type 4 secretion system (T4SS) represents a bacterial nanomachine capable of trans-cell wall transportation of proteins and DNA and which has attracted intense interest due to its roles in the pathogenesis of infectious diseases. During the current investigation we uncovered three distinct gene clusters in *Clostridioides difficile* strain 630 coding for proteins structurally related to components of the VirB4/D4 type 4C secretion system from *Streptococcus suis* strain 05ZYH33 and located within sequences of conjugative transposons (CTn). Phylogenic analysis shows that VirB4- and VirD4-like proteins of CTn4 locus, on one hand, and those of CTn2 and CTn5 loci, on the other hand, fit into separate clades, suggesting specific roles of identified secretion system variants in physiology of *C. difficile*. Our further study on VirB4- and VirD4-like products coded by CTn4 revealed that both proteins possess Mg^2+^-dependent ATPase activity, form oligomers (most probably, hexamers) in water solutions, and rely on potassium but not sodium ions for the highest catalytic rate. VirD4 binds nonspecifically to DNA and RNA. Its DNA binding activity strongly decreased with the W241A variant. Mutations in the nucleotide sequences coding for presumable Walker A and Walker B motifs decreased stability of the oligomers and significantly but not completely attenuated enzymatic activity of VirB4. In VirD4, substitutions of amino acid residues in the peptides reminiscent of Walker structural motifs resulted neither in attenuation of enzymatic activity of the protein nor influenced the oligomerization state of the ATPase.

**Importance:** *C. difficile* is a Gram-positive, anaerobic, spore-forming bacterium that causes life-threatening colitis in humans. Major virulence factors of the microorganism include toxins TcdA, TcdB and CDT. However, other bacterial products, including a type 4C secretion system, have been hypothesized to contribute to the pathogenesis of the infection and are considered as possible virulence factors of *C. difficile*. In the current paper we describe structural organization of putative T4SS machinery in *C. difficile* and characterize its VirB4- and VirD4-like components. Our studies, in addition to significance for basic science, can potentially aid development of anti-virulence drugs suitable for treatment of *C. difficile* infection.

## Introduction

*Clostridioides* (formerly *Clostridium*) *difficile* is a Gram-positive, anaerobic, spore-forming bacterium causing *C. difficile* infections (CDI) in humans, which include antibiotic-associated diarrhea, colitis, and pseudomembranous colitis (1). Studies on molecular pathogenesis of CDI demonstrated that the ability of the bacterium to cause the disease depends largely on the production of toxins – two large glucosylating toxins (TcdA and TcdB) and binary ADP-ribosylating toxin CDT (2-6). Deeper knowledge of the pathogenesis of CDI suggested that other microbial products in addition to the toxins can be of high value for the pathogenicity of the infection agent. Such non-toxin virulence factors can include surface structures of the bacterium participating in motility and adhesion (7-13).

Recently, components of a type 4 secretion system (T4SS) have been found in *C. difficile* genomes within the sequences of conjugative transposons (CTn) 2, CTn4 and CTn5 (14-16). Biological functionality of T4SS has not been directly demonstrated in *C. difficile*, although the transposons, containing T4SS components, were shown to excise from the genome of strain 630 and transfer to strain CD37 (17).

In general, the T4SS represents a nanomachine capable of transportation of proteins and DNA into eukaryotic target cells, horizontal interbacterial transfer of mobile genetic elements, and exchange of DNA with the outer space (18). Functionally, the T4SSs can be subdivided into two major classes. One class participates in interbacterial transfer of DNA through conjugation or by exporting/importing DNA to/from the extracellular space. It results in horizontal transfer of mobile genetic elements carrying beneficial for the microbial population traits. Another class is found in the Gram-negative bacteria like *Agrobacterium tumefaciens, Legionella*, and others; it is directly important for pathogenesis of the corresponding diseases and accomplishes its functions through delivery of effector DNA or proteins into target eukaryotic cell, thereby manipulating a plethora of eukaryotic functions (19, 20).

On a structural basis, the T4SS has been classified into types A, B and C. The type representatives of T4SS-A include the *A. tumefaciens* VirB/VirD4 T4SS and *E. coli* conjugation machinery of the R388 and pKM101 plasmids. In the former case, the T4SS-A is composed of 12 subunits, each in multiple copies, termed VirB1 through VirB11 and VirD4: (a) the cytoplasmic ATPases (VirB4, VirB11, VirD4), (b) components of an inner membrane complex (VirB3, VirB6, VirB8), (c) components of an outer membrane complex (VirB7, VirB9, VirB10), and (d) pilus components (VirB1, VirB2, VirB5). The prototype T4SS-B is the Dot/Icm secretion system of *L. pneumophila*. It is composed of 27 subunits, of which only few are structurally related to those of T4SS-A, while being functionally similar (18, 21, 22).

Recently, a new variant of T4SS (T4SS-C) was identified within the 89 kb pathogenicity island of *Streptococcus suis* (23). Only four components of this secretion system (VirB1, VirB4, VirB6, and VirD4) appeared to be sufficient for its function. As suggested by the authors, a VirB1-like sequence encodes an amidohydrolase with the CHAP domain, which is responsible for punching holes in the peptidoglycan; VirB6 probably forms the transportation channel across the cell wall, and VirB4 and VirD4 proteins energize the whole secretion device (24). Interestingly, the intact T4SS-C has been shown to be necessary for development of streptococcal toxic shock syndrome (23, 25, 26) and secretion of subtilisin-like proteinase and potential effector molecule SP1 (24, 27, 28).

Since components of T4SS occurring in pathogenic *C. difficile* isolates might play roles in the pathogenesis of CDI (14, 15), we aimed our study at characterization of VirB4- and VirD4-like proteins, elements of this newly identified secretion system.

## Results

### Characterization of VirB4/D4-coding loci in *C. difficile* 630 genome

As a first step toward characterization of VirD4 and VirB4 proteins, we looked into organization of T4SS loci in the genome of *C. difficile* 630 utilizing the data on T4SS-C of *S. suis* 05ZYH33 (24). Using amino acid sequences of VirB4 (NCBI database protein tag ABP89935.1) and VirD4 (ABP89939.1) of *S. suis* in a BLAST search (https://blast.ncbi.nlm.nih.gov/Blast.cgi), three loci containing similar proteins could be identified in *C. difficile* [Figure 1]. One locus was found on a complementary strand of chromosomal DNA within the nucleotide sequences of CTn4 (named therefore here “VirB4/D4_ CTn4”), while the two other were on a direct strand (i.e. “VirB4/D4_CTn2” and “VirB4/D4_CTn5”).

**Figure 1.**
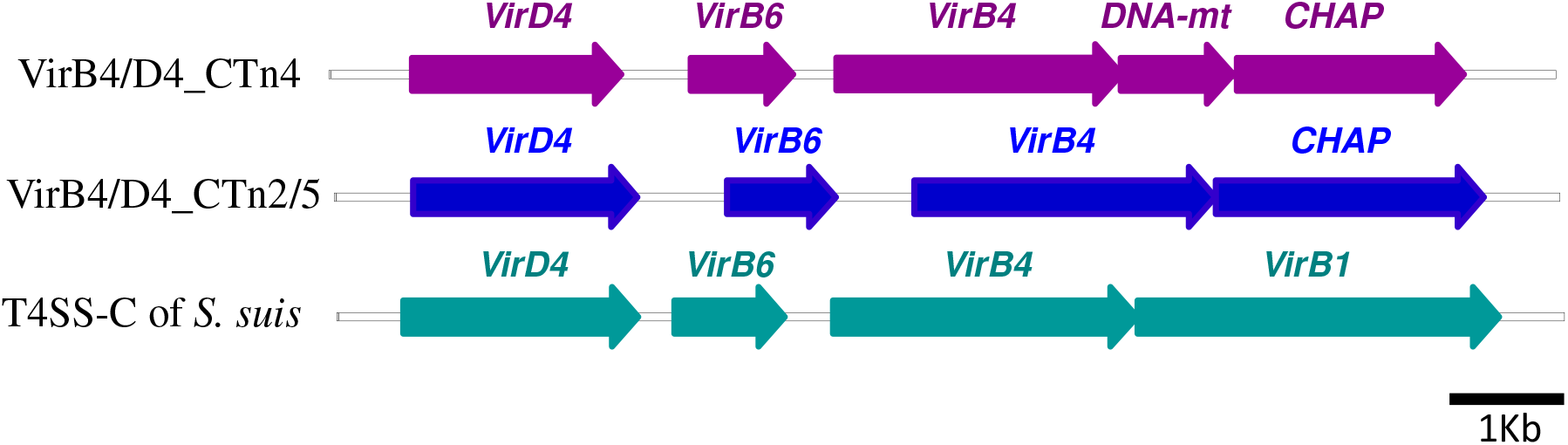
Genetic organization of VirB4/VirD4 loci in *C. difficile* 630. DNA-mt, DNA-methyltransferase; CHAP, CHAP domain-containing protein. The coding sequences are shown approximated to scale. See Table S1 for precise identification of the *C. difficile* 630 T4SS components.

The three VirB4/D4 loci consist of VirD4-, VirB6-, and VirB4-coding genes in an orientation identical to that of *S. suis*. The last coding sequence in the *S. suis* operon is *virB1*. In contrast, *C. difficile* contains here an unstudied open reading frame annotated as “CHAP domain-containing protein”. In the case of VirB4/D4_CTn4, there is an additional DNA-methyltransferase-coding sequence between the genes coding for VirB4 and CHAP domain-containing protein. VirB4 and VirD4 from CTn4 locus were 56-58% and 74% identical to the corresponding proteins coded by CTn2/5 loci, respectively, while identity of the matching proteins within CTn2/5 loci exceeded 90%. VirB4 and VirD4 of *S. suis* contained 31-36% of identical amino acid residues with VirB4_CTn2/4/5 or VirD4_CTn2/4/5 [Figure S1]. These data suggested that according to the primary sequences, the identified *C. difficile* proteins represented components of two putative secretion machineries: VirB4/D4_CTn4 and VirB4/D4_CTn2/5, each of which was only distantly related to described previously canonical T4SS-C apparatus from *S. suis* (24).

To get more information on relationship of *C. difficile* T4SS variants with known secretion systems, we used BLAST searches utilizing VirB4 and VirD4 protein sequences from CTn4 and CTn2/5 clusters as queries. Surprisingly, the database examinations revealed several groups of bacterial products which were over 90% identical to VirB4 and VirD4 from *C. difficile* and clearly different from the components of previously investigated T4SS-C from *S. suis* or *E. faecalis* [Figure S2]. Thus, CTn2/5 group contained VirB4/D4 components from pathogenic Gram-positive microorganisms like *Enterococci, Streptococci*, and *Peptostreptococci*, while CTn4 group included VirB4/D4 proteins from organisms of poorly studied genera *Blautia, Coprococcus*, and *Eisenbergiella*. Since we failed to find any data on T4SS organization of the latter bacteria, we focused our further investigations onto VirB4 and VirD4 components of CTn4.

### Purification of recombinant VirB4_CTn4 and VirD4_CTn4 proteins

According to the nucleotide sequence data, VirD4 (WP_011861117.1) is a 595-amino acid residues protein with a molecular mass of 68 kD, while VirB4 (WP_011861114.1) consists of 809 amino acid residues with a calculated mass of 91 kD.

The coding sequences were amplified using PCR and cloned into a pET28a vector. However, our attempts to produce the full-size proteins in *E. coli* strains transformed with the corresponding plasmids were unsuccessful. As seen in SDS-polyacrylamide gels, only minute amounts of the proteins could be isolated by HisTrap affinity chromatography, and the quality of the purified proteins was not sufficient for further studies [Figure S3]. Bearing in mind that the investigated proteins are products of a Gram-positive microorganism, we tried commercially available *Bacillus megaterium* expression system for production of VirD4 and VirB4. In contrast to VirD4, VirB4 could be isolated from *B. megaterium* in sufficiently pure form. However, the purified protein lacked any ATPase activity [Figure S4].

It is known that VirB4 and VirD4 consist of several structural parts, including COOH-terminal nucleotide-binding domain (NBD), which facilitates ATPase activity (29). Due to our special interest in enzymatic properties of the VirB4/D4 proteins, we performed NH2-terminal truncations by PCR and cloned the coding sequence of the remaining COOH-terminal NBDs into pET28 vectors. The engineered delVirB4 and delVirD4 consisted of amino acid residues 438-809 and 110-595, respectively, fused to 6xHis-containing NH2-terminal tag. Unfortunately, as with the full-size proteins, production of the deleted versions of the proteins in *E. coli* was also insufficient for biochemical analysis (data not shown).

To improve expression level of the delVirB4 and VirD4 proteins, we utilized MBP-fusion vector pMal-c5x. Addition of the normally well-produced hydrophilic MBP tag to the protein sequences considerably increased expression level of the coding sequences and kept the bacterial products in soluble form, thus helping isolation of relatively high amounts of the enzymes [Figure S3B]. Purification of delVirB4 and delVirD4 from liquid TB cultures resulted in a higher yield, as compared to LB, although the proteins appeared more degraded. This fact eliminated any advantages of the high protein production in the former medium. Thus, using pMal-based plasmids we were able to repeatedly isolate 3-4 mg of pure protein from 1 L of liquid LB culture. Therefore, in subsequent experiments deleted versions of VirB4 and VirD4 were used as MBP-tagged constructs.

### Enzymatic characterization of delVirB4_CTn4 and delVirD4_CTn4

Initial experiments with different concentrations of delVirB4 and delVirD4 demonstrated dose-response kinetics. In contrast to a negative control (MBP), increasing concentrations of *C. difficile* proteins resulted in an efficient hydrolysis of added ATP over a 1h-period at 35°C [Figure 2A, S5A]. Time curve data demonstrated an almost linear phosphate release curve over the period of 2 h for delVirB4, while delVirD4 appeared to be less stable at later time points under the conditions of the experiments [Figure 2B]. Both proteins were active at temperatures of 23°C – 42°C [Figures 2C, S5B]. Higher temperature appeared to be detrimental for delVirD4, while delVirB4 resisted incubation at 42°C for 30 min quite well. Lastly, the two proteins demonstrated obvious preference for ATP over GTP [Figure 2D].

**Figure 2.**
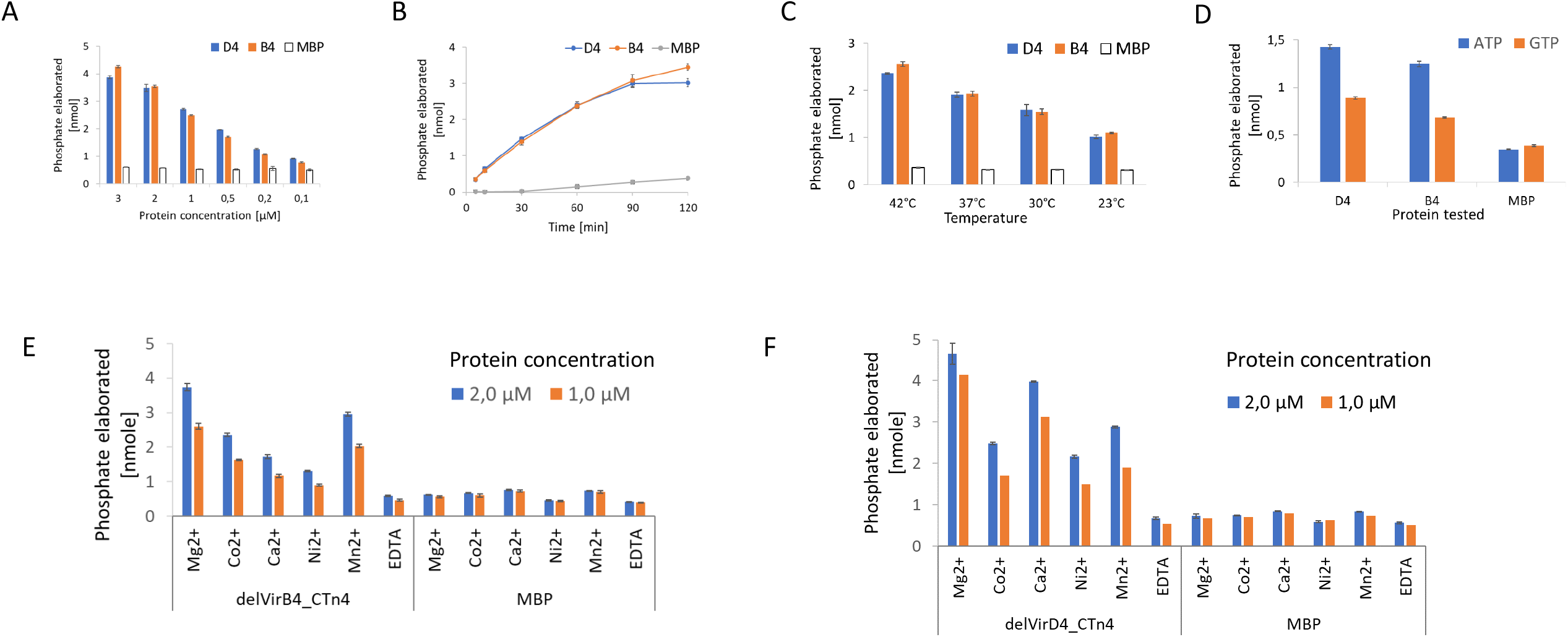
Influence of multiple parameters on NTPase activity of delVirD4_CTn4 (D4) and delVirB4_CTn4 (B4). Panel A, the proteins at indicated concentrations were incubated at 35°C with 100 µM ATP for 1h. Panel B, the proteins at 1 µM were incubated with 100 µM ATP for indicated time. Panel C, the proteins at 1 µM were incubated with 100 µM ATP at indicated temperatures for 30 min. Panel D, the proteins at 1 µM were incubated at 35°C for 1h with 50 µM ATP or GTP. Panels E and F, Indicated cations (used as chloride salts) or EDTA (each 2 mM) were added to the reaction mixtures, containing 100 µM ATP and 1 µM or 2 µM of delVirB4_CTn4 (Panel E) or delVirD4_CTn4 (Panel F), and incubated at 35C for 1 h.

In the next set of experiments, we studied influence of cations on enzymatic performance of delVirB4 and delVirD4. Addition of EDTA completely blocked the reaction. The highest ATPase activity was seen with MgCl2 for both proteins, followed by MnCl2 and CaCl2 with delVirB4 and delVirD4, respectively [Figure 2E, F]. When potassium was used as a substitute for sodium ion in a reaction mix, it significantly stimulated ATPase activity of both proteins, as clearly seen from the kinetic data [Figure 3].

**Figure 3.**
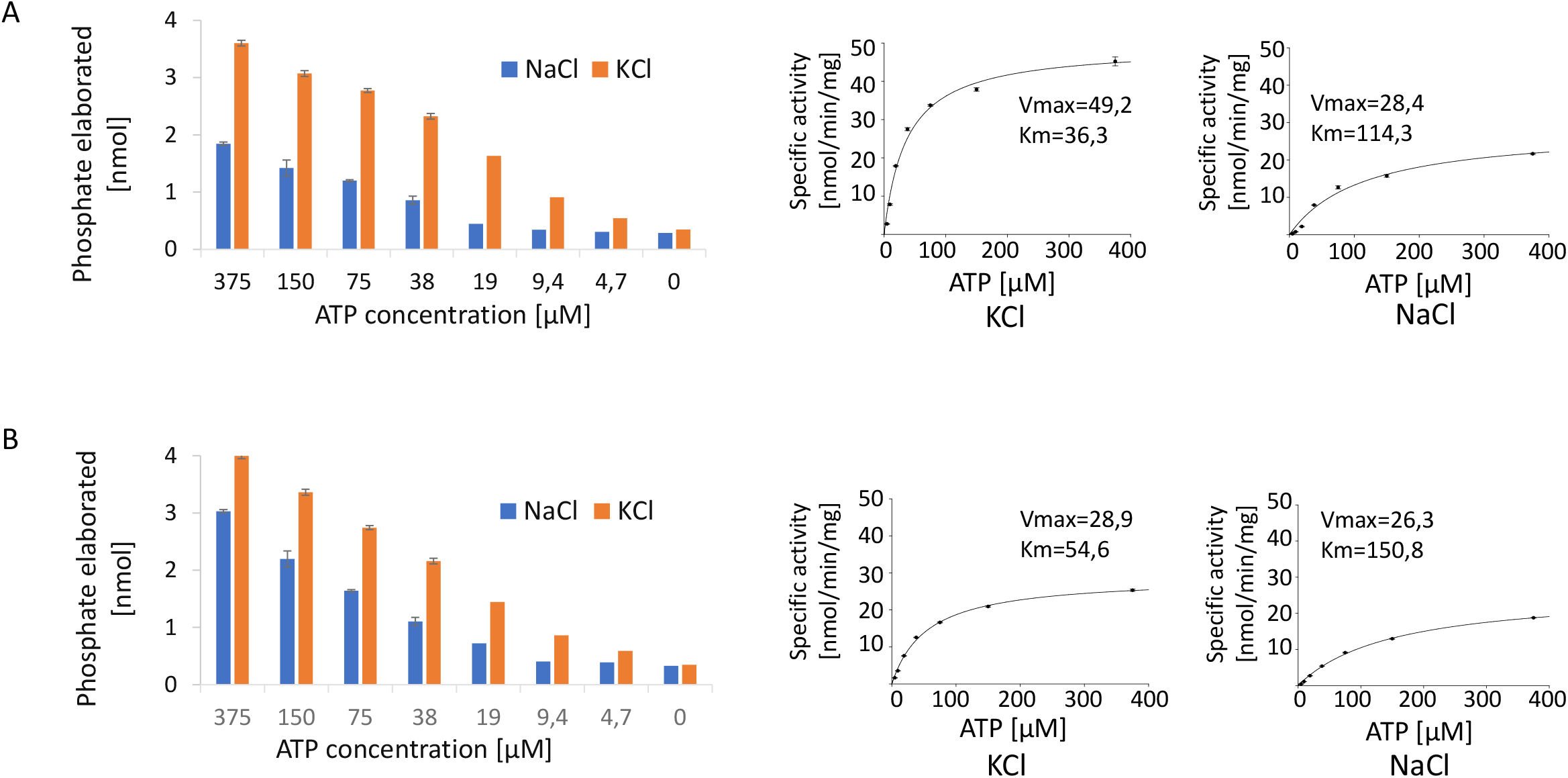
Dependance of ATPase activity on potassium vs. sodium cation. DelVirB4_CTn4 (Panel A) and delVirD4_C (Panel B) at 0,25 µM were incubated for 1 h at 35°C with different concentrations of ATP in Tris-HCl buffer containing 2 mM MgCl_2_ and 150 mM NaCl or 75 mM KCl.

### Oligomer formation by delVirB4_CTn4 and delVirD4_CTn4

To investigate the stoichiometry of delVirB4 and delVirD4 under native conditions, we used gel chromatography. Concentrated preparations of the ATPases were loaded onto Superdex200 column and eluted with TBS. Surprisingly, no protein peaks were seen at expected elution volumes (∼14 ml for proteins ∼100 kD). Instead, two peaks could be observed at elution volumes of ∼8 ml (which is close to the void volume of the column) and around 15 ml (where proteins of 40-50 kD should elute) [Figure 4A]. SDS-PAGE analysis showed that material eluted near the void volume contained delVirB4 and delVirD4, while in 15-ml material fragments of 40-50 kD could be seen [Figure 4BC]. These data indicated that in water solutions both proteins formed high-molecular mass oligomers. Bearing in mind technical characteristics of the Superdex200 10/30 column, the molecular mass of the protein complexes should exceed 600 kD, suggesting oligomer formation with a stoichiometry of ≥ 6 monomers.

**Figure 4.**
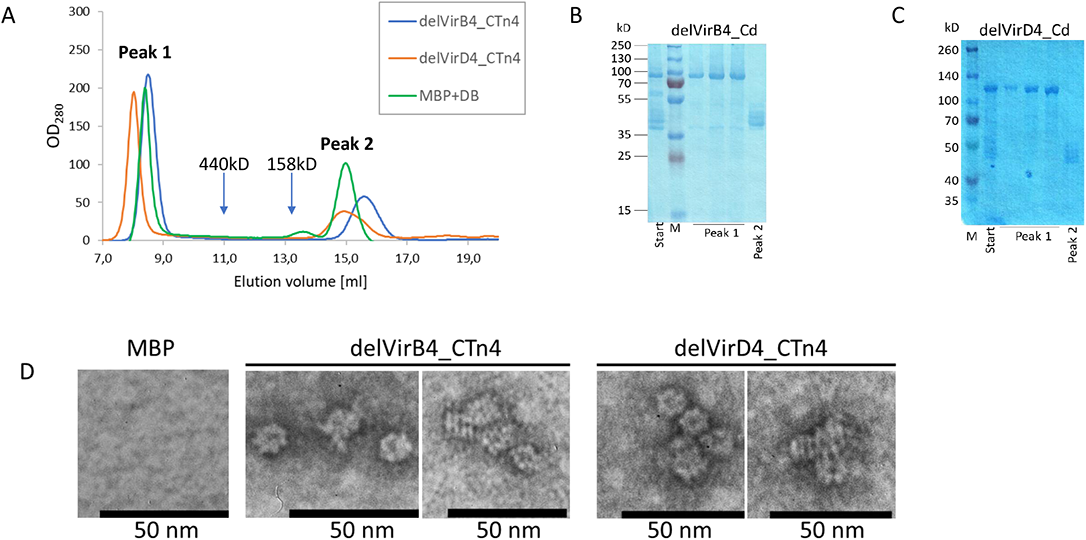
Oligomerization of MBP-tagged delVirB4_CTn4 and delVirD4_CTn4. Panel A, overlay of partial Superdex200 chromatograms with delVirB4_CTn4, delVirD4_CTn4, and a mixture of MBP (45kD) plus Dextran Blue (DB, 2MD) as size markers. Approximate positions of ferritin, 440kD and aldolase, 158 kD are indicated by the arrows. Panels B, and C, SDS-PAGE analysis of MBP-tagged delVirB4_CTn4 (Panel B), and delVirD4_CTn4 (Panel C) loaded on (Start) and eluted from (Peaks 1 and 2) Superdex200 column. Lane M, molecular mass of markers is shown in kilodaltons (kD) on the left. E. Proteins after MBPtrap chromatography were additionally purified by Superdex200 (see analysis of peak 1 fractions on panels B, and C), diluted till 10 µg/ml with dH_2_O, and subjected to negative staining/transmission electron microscopy. Samples were processes as described in Materials and Methods. Black horizontal scale bar represents size of 50 nm.

To further study oligomerization processes of delVirB4 and delVirD4 proteins, we used method of electron microscopy with negative staining. During these experiments, we saw particles of ∼10 nm, demonstrating predominantly two types of silhouettes. Most of them were circular and exhibited 6-7-fold symmetry, while others have a striated rectangular look [Figure 4D]. Such structural variability probably resulted from different plane positions during sample processing in electron microscopy.

### Structure-function analysis of delVirB4_CTn4 and delVirD4_CTn4B

To probe the structure-function relationship in delVirB4 and delVirD4, we changed amino acid residues within the regions known to be important for catalytic activity of the ATPases. In relation to delVirB4, these include G420 and K421, belonging to a Walker A motif, and D633 and E634 of Walker B. With delVirD4, the substituted residues were K152 and T153 of Walker A, and D408 and E409 in Walker B motifs [Figure S1]. In accordance with the literature (30), all engineered mutations decreased ATPase activities of the VirB4 protein. Interestingly, increasing the negative net charge in the Walker A motif of delVirB4 by introducing D420 or E421 produced less of an effect than changing the side chain structure in G421 variant. Mutations in the Walker B-coding sequence (D633K and E634K), resulting in inversion of charge in the corresponding sites, appeared to suppress the ATPase activities of delVirB4 more prominently than mutations of Walker A [Figure 5AB]. Mutagenesis experiments performed on VirD4 failed to demonstrate any involvement of the substituted amino acid residues in catalysis.

**Figure 5.**
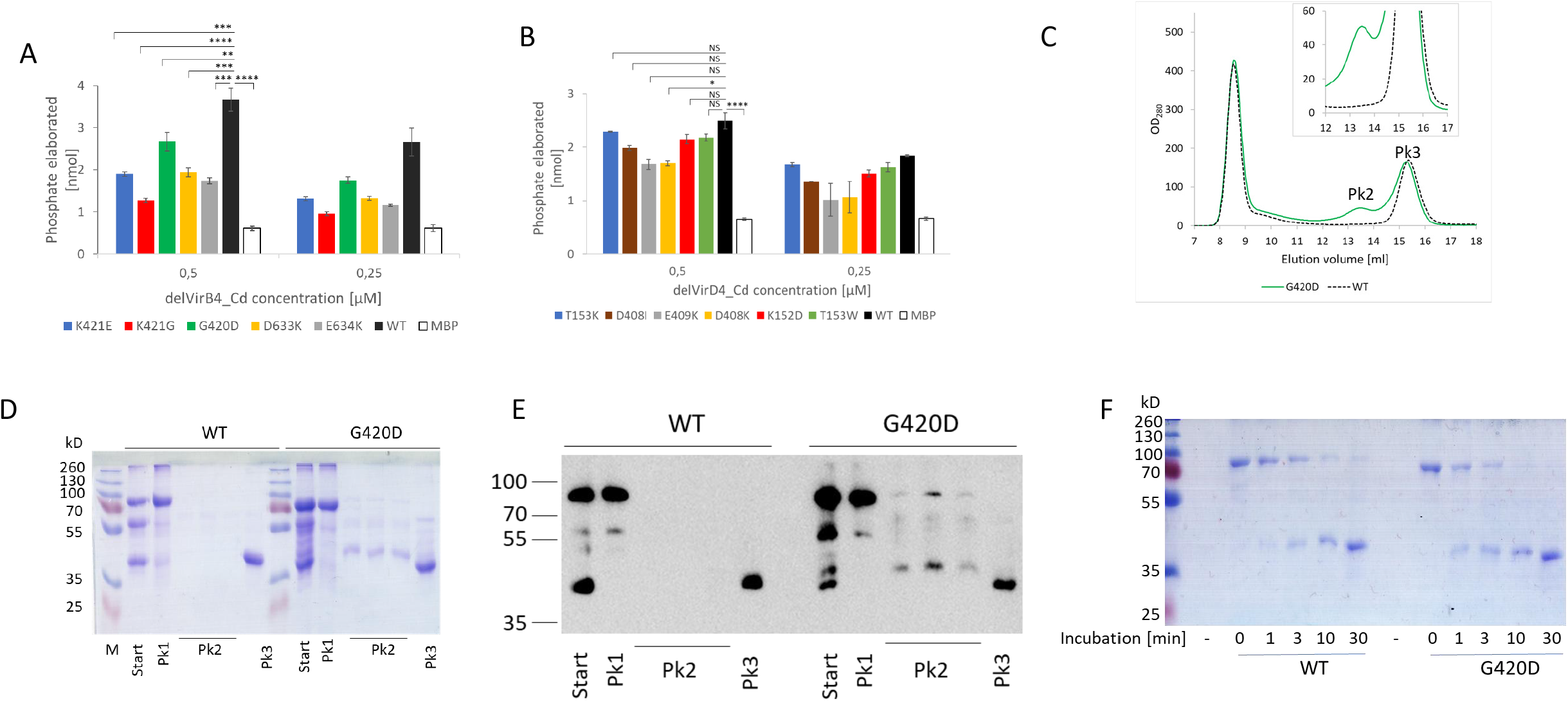
ATPase activity and oligomer stability of site-substituted delVirB4_CTn4 and delVirD4_CTn4. Panels A and B, indicated amino acid residues were substituted by QuikChange mutagenesis on the corresponding wild type coding sequences. Enzymatic reaction was performed with two protein concentrations (0,5 and 0,25 µM) and 150 µM ATP. Results are shown as means of 2 independent experiments with 3 measurement replicates each. Error bars represent standard deviations (n=6). Student’s t test was applied for statistical comparisons. **p<0,05; ***p < 0.001; ****, p<0,0005; NS, not significant. Panels C – E, oligomer stability of the wild type and G420D delVirB4_CTn4 variants. Purified by MBP-tag chromatography delVirB4_CTn4 variants we subjected by Superdex200 chromatography in 20mM Tris-HCl, pH=7,4 plus 75 mM KCl (Panel C). Thereafter, starting material (Start) and that of the major peaks (Pk 1, 2, and 3) were analyzed in 10% SDS-PAGE (Panel D) and Western blotting with anti-MBP serum (Panel E). Fractions of peak 1 obtained from WT and G420D VirB4 variants were pooled and subjected to analytical trypsinolysis (1 ng of trypsin per 2 µg of the protein) for 1, 3, 10 and 30 min at 25°C and SDS-PAGE analysis (Panel F).

In the next set of experiments, we compared oligomer structure stability of the wild type delVirB4/D4 proteins with that of the site mutants. Using Superdex200 gel-chromatography, SDS-PAGE, Western blotting, and trypsin cleavage experiments we demonstrated that all amino acid substitutions in delVirB4 but not in delVirD4 variants resulted in oligomer instability, manifested by the appearance of an additional peak on the chromatograms [Figure 5, S6]. This peak contained monomeric full size delVirB4 and its fragments [Figure 5C-E]. Although the majority of the molecules remained in an oligomeric state and constituted the material of peak 1, trypsinization experiments verified its higher susceptibility to cleavage in the tested G420D variant in contrast to the wild-type ATPase [Figure 5F].

### Nucleic acid-binding activity of delVirB4_CTn4 and delVirD4_CTn4

In the next set of experiments, we studied interaction of the *C. difficile* ATPases with DNA and RNA. To this end, following incubation on ice for 30 min, mixtures of delVirB4 or delVirD4 with the plasmid dsDNA, chromosomal dsDNA, or rRNA were separated by 0.5% agarose gel electrophoresis. In contrast to delVirD4, delVirB4 did not bind dsDNA. delVirD4 demonstrated stable dsDNA-protein and rRNA-protein interactions, illustrated by gel shift of ethidium bromide-stained DNA and RNA fragments [Figure 6]. Chelation of cations by EDTA significantly decreased DNA-binding activity of delVirD4, while addition of divalent salts restored complex formation [Figure 6D, S7]. To test if ssDNA is also able to interact with delVirD4, we performed competition experiments, in which preformed protein-plasmid complex was treated with oligonucleotide ssDNA. Addition of the 81-mer oligonucleotide primer in increasing amounts displaced dsDNA from the preformed complex, which was evidenced by the appearance of a ∼ 5 kb plasmid band [Figure 6E]. With the plasmid, complex formation was independent on the form (linear or circular) of the added DNA molecule [Figure 6F]. Site-substitution W241A in a presumable nucleic acid-binding region (31) resulted in a strong decrease in DNA binding by delVirD4 [Figure 6G]. Interestingly, the mutation influenced neither ATPase activity of the protein nor its oligomerization state [Figure S6B, S8].

**Figure 6.**
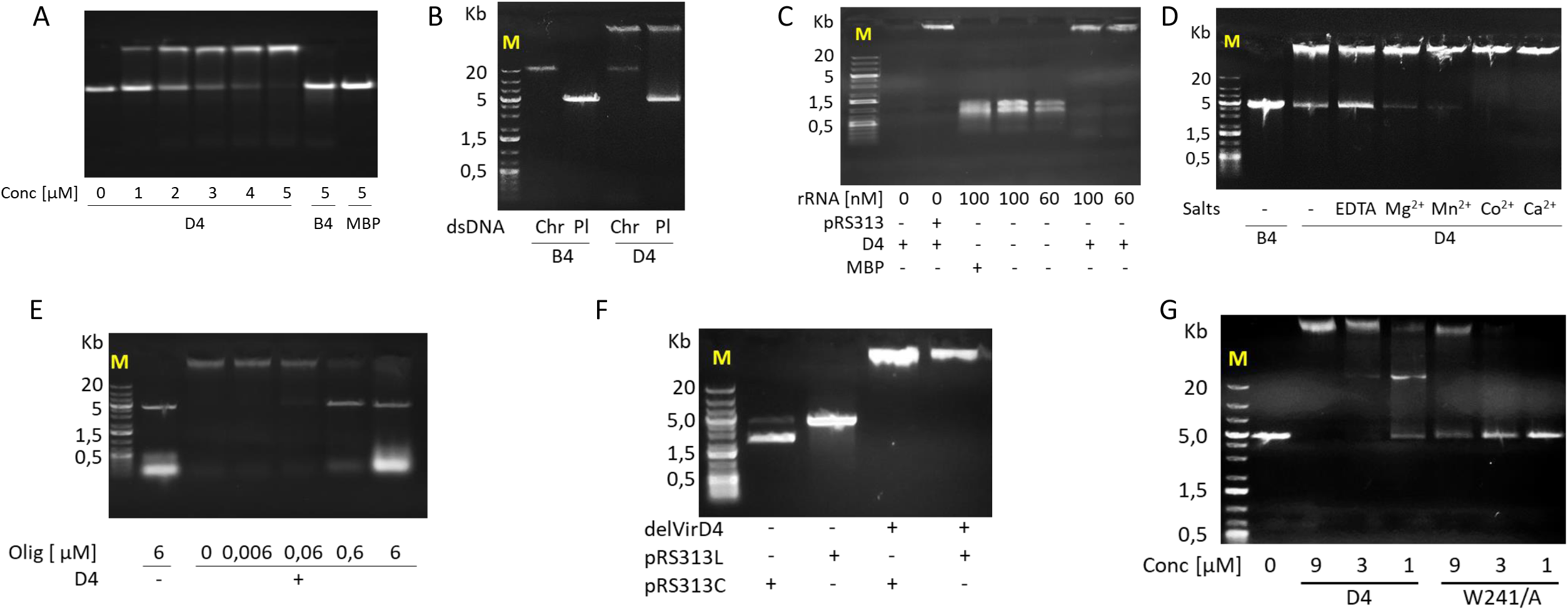
Interaction of delVirB4_CTn4 (B4), wild type delVirD4_CTn4 (D4) and W241A variant of delVirD4 (W241A) with nucleic acids. A. Binding of plasmid dsDNA (pRS313, 3,3 nM) with the *C. difficile* proteins at different concentration. B, binding of the *C. difficile* proteins (10 µM) with plasmid (Pl, 3,3 nM) and chromosomal (Chr, 50 ng) dsDNA. C, Binding of the delVirD4_CTn4 protein (D4) with rRNA. RNA at indicated concentrations (or pRS313 at 3,3 nM as a positive control) was mixed with 10 µM delVirD4_CTn4 (or MBP) and processed as described in Materials and methods. D, Binding of the *C. difficile* proteins (3 µM) with plasmid dsDNA (pRS313, 3,3 nM) in the presence of added EDTA or metals salts (2 mM each). Addition of EDTA or metal cations to plasmid DNA without *C. difficile* proteins did not change nucleic acid mobility patterns [Figure S6]. E, interaction of delVirD4_CTn4 with synthetic DNA oligonucleotide ssDNA. delVirD4_CTn4 (10 µM) was incubated with pRS313 (2nM) for 30 min on ice. Thereafter, indicated amounts of oligonucleotide ssDNA were added and incubation on ice continued for 30 min. F, Plasmid pRS313 was linearized by BamHI/EcoRI cleavage or left untreated as circular. Thereafter, linear (pRS313L) and circular (pRS313C) plasmid dsDNA (3,3 nM each) were mixed with delVirD4_CTn4 *C. difficile* protein (10 µM) and incubated for 30 min on ice. Agarose gel electrophoresis was performed in 0,5% agarose (A, B, C, D, F) and 1,5% (E). G, influence of mutations on Interaction of delVirD4_CTn4 with plasmid DNA. Wild type and mutated delVirD4_CTn4 at indicated concentrations were incubated with the plasmid pRS313 (2,5nM) for 30 min on ice. Thereafter, the samples were subjected to electrophoresis in 0,5% agarose. M, nucleic acid marker. Size of major fragments is shown on the left in Kb.

Due to the fact that binding of DNA can affect the enzymatic properties of *C. difficile* proteins, we studied influence of DNA on ATPase activity of delVirB4 and delVirD4. It was not obviously the case, given that the addition of pRS313 (dsDNA) or synthetic oligonucleotide (ssDNA) to the ATPase reaction mix did not change enzymatic activity of the proteins [Figure S9].

## Discussion

Nine types of microbial secretion systems are known to date, among which type 4, comprising type 4A and type 4B subfamilies, is relatively well studied and is able to deliver its substrates (DNA or proteins) to eukaryotic or prokaryotic cells (19, 32). Recently, subfamily T4SS-C was found in *C. difficile* (14, 15), though neither its biological functionality nor biochemical properties of its components have been studied.

Genes coding for VirD4, VirB4, and VirB6 can be identified in *C. difficile* 630. However, instead of VirB1 coding sequence present in *S. suis*, a “CHAP domain-containing protein” is encoded in *C. difficile* chromosome as a last gene of the operon. “CHAP” stands for <u>c</u>ysteine, <u>h</u>istidine-dependent <u>a</u>midohydrolases/<u>p</u>eptidase and is generally involved in partial degrading cell wall of Gram-positive microorganisms during cell division or phage release (33, 34). On the other hand, a CHAP-related sequence can be found as a domain in a VirB1-like protein of *S. suis* (24). Based on these data, one can speculate that the coding sequence, termed CHAP domain containing protein, can accomplish the functions of VirB1 in *C. difficile* and is involved in local degradation of clostridial cell wall during T4SS channel assembling.

An interesting feature of VirB4/D4_CTn4 operon is the presence of a DNA methyltransferase coding sequence. There are three known major forms of DNA methylation in prokaryotes, accomplished by DNA methyltransferases: 6-methyladenine, 4-methylcytosine, and 5-methylcytosine. Increasing evidence suggests that DNA methylation regulates a number of biological processes of high medical importance, including antibiotic resistance, virulence, evasion of host immune response, and replication niche adaptation (35-38). In a recent paper, conserved 6-methyladenine methyltransferase (*camA*) was identified as important factor for sporulation of *C. difficile* and pathogenesis of experimental CDI (39). Whether a DNA methyltransferase encoded by the VirB4/D4_CTn4 locus is involved in the pathogenicity of *C. difficile* is unknown, as is the relationship of the DNA methyltransferase with the T4SS-C function.

The major components of T4SSs include two VirB4- and VirD4-related ATPases. As seen from our protein homology searches, VirB4_CTn4 and VirB4_CTn2/5 fall into two different clades, as do VirD4_CTn4 and VirD4_CTn2/5, which, in turn, are distinct from the clade containing previously described “classical” T4SS-C of *S. suis* (24). Intriguingly, some strains of *S. suis* possess an operon related to VirB4/D4_CTn2/5 of *C. difficile* instead of “classical” T4SS-C. These data illuminate a fruitful direction of further research on the structure-functional variability in type 4 secretion machinery of Gram-positive microorganisms.

Most biochemical data on VirB4 and VirD4 were obtained with the proteins originated from Gram-negative microorganisms, predominantly, from *E. coli* (40-43). Bearing in mind the low sequence similarity of the clostridial ATPases with those of *E. coli* T4SS, it was interesting to study catalytic properties of VirB4 and VirD4 homologs from Gram-positive human pathogen *C. difficile*. We found that both delVirB4_CTn4 and delVirD4_CTn4 preferred higher temperatures for enzymatic activity, up to 42°C. This was in some contrast to previous data on *E. coli* VirD4 homolog TrwB that demonstrated an optimal temperature range of 33-37°C, with a drastic drop at 42°C or higher (31). Similar to TraE and TraK (VirB4 and VirD4 homologs in *Aeromonas veronii* T4SS, respectively), the best cofactor in the reaction with the clostridial proteins was Mg^2+^. However, the second most efficient cations with delVirB4_CTn4 and delVirD4_CTn4 were Mn^2+^ and Ca^2+^, while for TraE and TraK these were Mn^2+^ and Co^2+^ (44). In line with the previous data on T4SS ATPases, sodium ions were strongly inhibitory for *C. difficile* VirB4/D4 enzymes and should be replaced in the reaction by potassium (40).

Several studies with purified VirB4/TraB, VirB4/TrwK, and VirD4/TrwB of *E. coli* demonstrated that the proteins in water solution behaved as enzymatically active hexamers (40, 42, 43, 45). These data were confirmed by gel filtration, sedimentation velocity analysis, native polyacrylamide gel electrophoresis, and electron microscopy. However, in other experiments, purified VirB4 behaved as monomers (29, 30). In our experiments both delVirB4_CTn4 and delVirD4_CTn4 formed high-molecular mass oligomers and hexameric particles, as seen in gel chromatography and electron microscopy, respectively. Moreover, as observed during Superdex200 chromatography, the oligomerization rate for both proteins was close to 100% since no traces of monomers or dimers could be found on chromatograms.

Two peptides are known to represent general nucleotide-binding motifs of ATP-requiring enzymes: Walker A (or P loop) and Walker B. The consensus sequence of Walker A was originally identified as GxxxxGKT/SxxxxxxI/V, while Walker B conserved features are represented by R/KxxxGxxxLhhhhDE (“h” stands for a hydrophobic amino acid residue) (46). According to crystal structure of the VirD4/TrwB hexamer, Walker A and Walker B are located on the surface of each oligomer on the interface between the neighboring subunits. Walker A is preceded by a β-strand, followed by an α-helix, and forms a loop around bound ATP. Amino acid residues of the motif interact with phosphates of the nucleotide and Mg^2+^ (47). It is thus not surprising that amino acid substitutions in Walker A resulted in drastic drop of both ATPase and conjugating-stimulating activities (31, 48). Putative Walker A and Walker B motifs are easily detected in the VirB4_CTn4 sequence. We performed site-specific mutagenesis experiments and engineered a panel of amino acid substitutions to study their influence on ATPase activity of the proteins. All engineered proteins demonstrated significantly lower but still detectable levels of enzymatic activity, suggesting presence of cryptic catalytic centers in the proteins. Indeed, several conserved sequences marking ATP-processing proteins are known. In the case of VirB4, these include regions C (Dx[D/E]xxxE), D (RK), and E (S/TQ) (29). Additionally, a cryptic but functional novel ATP-binding site has been identified in the N-terminal domain of VirB4 (42). We did not perform any site-mutational experiments on these zones of *C. difficile* ATPases. Interestingly, attenuation of ATPase activity paralleled to less efficient oligomer formation as observed in our gel-chromatography and trypsinization experiments. Due to this observation, we are tempted to speculate that diminished enzymatic activities in the mutated VirB4 variants were caused primarily by structural perturbations in the molecules resulting in less efficient formation of functionally competent oligomers.

In contrast to VirB4, Walker A and Walker B sequences are not easily seen within the VirD4_CTn4 sequence. While we identified peptides, which can represent the motifs, and performed a set of mutation experiments, we are not absolutely sure in the correct identification of Walker A and Walker B positions in this case, since the results of our enzymatic experiments were negative.

Since VirB4/D4 T4SSs are known to participate in transporting DNA, we were interested if VirB4 and VirD4 ATPases of *C. difficile* could bind nucleic acids. According to our gel-shift experiments, delVirB4_CTn4 but not delVirB4_CTn4 formed complexes with DNA and RNA. In relation to VirD4, our results are in line with the published data. Indeed, several VirD4-like proteins have been shown to interact with both dsDNA and ssDNA (48-51). With VirB4 the data is controversial. VirB4/TraB, the ATPase of *E. coli* conjugation plasmid pKM101, formed complexes with DNA as demonstrated by agarose gel-shift and proteolysis protection assays (42), while VirB4/TrbE of the pRP4 plasmid and VirB4/TrwK of R388 did not (30). We are not aware of any published data showing interaction of VirD4 ATPase with RNA. Our experiments did demonstrate this binding behavior. It is interesting to speculate if RNA could be a T4SS substrate, and whether RNA translocation is important in some way for physiology or pathogenicity of the microorganisms.

Previously, it has been shown that replacement of W216, located in the central pore of the VirD4/TrwB *E. coli* hexamer, resulted in a protein that did not hydrolyze ATP and exerted a dominant negative effect on R388 conjugation (31). These tryptophan residues from adjacent monomers form a ring, which possibly participates in DNA binding and translocation. In our experiments, substitution of similarly located amino acid residue W241 by alanine produced a protein with wild type level of ATPase activity, normal oligomerization properties but diminished DNA-binding activity.

In summary, we identified novel variants of T4SS-C in important human pathogen *C. difficile* and characterized biochemical propertied of major VirB4- and VirD4-like ATPases. Our findings expand the list of bacterial secretion systems potentially important for bacterial physiology and pathogenicity and should be accompanied in future by the demonstration of T4SS functionality in *C. difficile*.

## Materials and methods

### Materials and bacterial strains

Restriction endonucleases, T4 DNA ligase, Phusion DNA polymerase, molecular mass markers and kits for DNA isolation were from Thermo Fisher Scientific (Moscow, Russia). Lysogeny broth Miller recipe (LB) medium was from Amresco (Solon, OH, USA), terrific broth (TB) medium, brain heart infusion and yeast extract were from Difco (Franklin Lakes, NJ, USA), liquid chromatography media were from GE Healthcare (Moscow, Russia), reagents for agarose, polyacrylamide gel electrophoresis and general laboratory reagents were from Merck (Moscow, Russia).

Molecular cloning and recombinant protein production were performed in *Escherichia coli* DH10B, Rosetta (DE3) (Merck) and *Bacillus megaterium* WH320 (MoBiTec GmbH, Goettingen, Germany). Plasmids for cloning and recombinant protein expression in *E. coli* were based on pUC19 (New England Biolabs), pET28a-c (cloning with NH2-terminal 6His tag, Merck), and pMal-c5x (cloning with the NH2-terminal maltose-binding protein (MBP) tag, New England Biolabs). A vector pHis1522 (MoBiTec) was used for expression in *B. megaterium. C. difficile* 630 (TcdA^+^/TcdB^+^/CDT^+^ human isolate (16)) was used as a source of DNA for gene cloning. Anti-MBP antibody was from New England Biolabs (#E8032S).

### Gene cloning

For cloning VirD4_CTn4 and VirB4_CTn4 coding sequences, the corresponding genes were PCR amplified from genomic DNA of *C. difficile* with the primers #808/#809 and #812/#813 respectively (primer information is presented in Table S2). The synthesized sequences were cut with NdeI and EcoRI restriction endonucleases and ligated into similarly digested pET28a vector, producing plasmids p28-VirD4_CTn4 and p28-VirB4_CTn4. To make 5’-terminal truncations of VirD4_CTn4 and VirB4_CTn4 coding sequences, plasmids p28-VirD4_CTn4 and p28-VirB4_CTn4 were used in PCR as matrix DNA with the primers #1014/#17 and #1038/#17. The resulting amplified nucleotide sequences were digested with BamHI and EcoRI endonucleases and ligated into pET28b and pET28a to yield p28-delVirD4_CTn4 and p28-delVirB4_CTn4, respectively. For site-directed mutagenesis of putative Walker A and Walker B motifs, gene fragment from p28-delVirB4_CTn4 was cloned into pUC19 using BamHI/EcoRI restriction endonuclease sites. Following QuikChange reaction (Stratagene, Moscow, Russia) with the corresponding primers (see Table S2), mutated sequences were re-introduced back into pET28. QuikChange mutagenesis of VirD4 was performed directly on p28-delVirD4_CTn4 plasmid. In few cases with VirD4 where QuikChange protocol did not result in successful nucleotide substitutions, PCR splicing reactions have been accomplished (52). To produce coding sequences with the additional MBP-containing tag, the plasmids p28-delVirD4_CTn4 and p28-delVirB4_CTn4 as well as their site-mutated variants were digested with NcoI/SalI and the isolated insert sequences were ligated into pMal-C5x. The resulting plasmids were named pMal-delVirD4_CTn4 and pMal-delVirB4_CTn4 for the wild type genes. Indications of the engineered amino acid substitution for the corresponding protein variants were added to the plasmid name where applicable.

To engineer plasmids for expression in *B. megaterium*, full size VirB4_CTn4 and VirD4_CTn4 were amplified from p28-VirB4_CTn4 and p28-VirD4_CTn4 using primers #474/#475 and ligated into SacI/KpnI restriction endonuclease sites of pHis1522, producing pHis1522-VirB4_CTn4 and pHis1522-VirD4_CTn4.

### Recombinant protein purification

For recombinant protein production, *E. coli* Rosetta strain was grown in LB medium supplemented with the corresponding antibiotics (ampicillin at 100 µg/ml for pMal-c5x or kanamycin at 50 µg/ml for pET28 plasmids) until OD600=0.5. Induction of expression was performed overnight with 0.5 mM isopropyl β-D-1-thiogalactopyranoside (IPTG) at 22°C or for 1 hr with 1 mM IPTG at 37°C with pET28 or pMal-c5X-based plasmids, respectively. For production of proteins in Gram-positive host, recombinant *B. megaterium* were grown in 3.7% brain heart infusion+0.5% yeast extract liquid medium with tetracycline at 10 µg/ml till OD600=0.3. Induction of expression was performed with 0.5% xylose overnight at 22°C. Both *E. coli* and *B. megaterium* were lysed by sonication. The resulting extracts were subjected to centrifugation at 30000 g for 30 min to separate cytosolic (water-soluble) and membrane (water-insoluble but 6M urea-soluble) fractions. Recombinant proteins were subsequently purified via affinity chromatography using HisTrap or MBPtrap columns connected to an ÄKTA Explorer liquid chromatography system (GE Healthcare) according to the instructions of the manufacturer. The purified proteins were stored in 10% glycerol/20 mM Tris-HCl, pH=7.4, 150 mM NaCl (TBS) at -20°C.

### Determination of ATPase activity

ATPase activity of the purified proteins was determined using two kits – Enliten ATPase assay (FF2000, Promega, Moscow, Russia) and malachite green phosphate assay (MAK307, Merck). Both assays were performed in two steps as detailed in the corresponding user manuals. Typically, reaction mixture at the first step consisted of 0.1 mM ATP, 1 µM protein of interest, 2 mM MgCl2, 20 mM Tris-HCl, pH=7.4, 150 mM NaCl in a total volume of 30 µl or 80 µl for Enliten and malachite green assays, respectively. The reaction proceeded for 1 h at 35°C. Thereafter, at the second step, 30 µl of luciferin/luciferase reagent or 20 µl of malachite green reagent were added. The results were read on GloMax-Multi+ plate reader immediately for Enliten or following incubation at 22°C for 30 min for malachite green assay. With Enliten assay, intensity of the emitted light, following the addition of luciferin/luciferase reagent, is proportional to ATP concentration. Thus, measurement of the light intensity using a luminometer permits direct quantitation of non-degraded ATP, present in reaction mix. For malachite green assay, the intensity of the developing green color is proportional to the concentration of phosphate, elaborated during ATP hydrolysis, and was measured at 600 nm. Here, to convert optical density displayed as “optical unit” (OU600) into the amounts of degraded ATP, a calibration curve was constructed by plotting OUs obtained with phosphate standards, against the amounts of added phosphate.

### Study of oligomer formation by delVirB4_CTn4 and delVirD4_CTn4

Oligomerization of delVirB4_CTn4 and delVirD4_CTn4 proteins was studied by gel chromatography on Superdex200 10/30 GL column connected to ÄKTA Explorer liquid chromatography system (GE Healthcare). The proteins, dissolved TBS at concentrations of ∼ 1.5 mg/ml, were loaded onto the column equilibrated in TBS or 20 mM Tris-HCl, pH=7.4, 75 mM KCl (TBK) and were eluted at 0.5 ml/min. Dextran blue, ferritin, aldolase, MBP, bovine serum albumin, and ovalbumin) were used as molecular mass standards.

The ultrastructure of delVirB4_CTn4 and delVirD4_CTn4 oligomers was studied using a negative staining method with a transmission electron microscope JEM 2100Plus (JEOL, Japan). A drop of the purified proteins at concentration of 0.1-0.01 mg/ml in distilled water was placed on a formvar copper grid (SPI Supplies, USA) for 60 s. The excess of solution was removed by filter paper. A copper grids with the samples were stained with 1% uranyl acetate (Serva, Heidelberg, Germany) for 60 s, and the excess of stain was then removed by filter paper. Grids were dried at room temperature for 10 min and analyzed using transmission electron microscope at an accelerating voltage of 160 kV.

### Determination of DNA/RNA binding activity

To determine double-stranded (ds) DNA-binding activity, the purified NH2-terminal deleted MBP-tagged proteins in different concentration were mixed with 50 ng of *C. difficile* chromosomal DNA or with 3.3 nM of pRS313 plasmid DNA (53) linearized with BamHI/EcoRI restriction endonucleases. The mixtures were kept on ice for 10 min and then were subjected to 0.5% agarose gel electrophoresis in 40 mM Tris-20mM acetate-1 mM EDTA (TAE) buffer, stained with ethidium bromide. Interaction of a single-stranded (ss) DNA with delVirD4_CTn4 was studied in competition experiments, in which initially formed dsDNA-protein complex of 2 nM of pRS313 and 10 µM of the *C. difficile* protein was treated with increasing amounts (0.006 - 6 µM) of an 81-mer DNA oligonucleotide on ice for 30 min. Thereafter, the reaction was visualized by 1.5% agarose gel electrophoresis as described above. For studying interaction of ribonucleic acid with VirD4_CTn4 we used RNA, isolated from yeast ribosomes (a general gift of prof. Sabine Rospert, Freiburg University, Germany (54)) by RNeasy kit (Thermo Fischer Scientific). Purified RNA at concentrations of 60 nM and 100 nM was added to the indicated proteins and processed as described above for dsDNA.

### General biochemical methods

Purified protein preparations were analyzed by polyacrylamide gel electrophoresis (PAGE) in sodium dodecyl sulfate (SDS)-containing buffer (55) and Western blotting with the anti-MBP antibody (56). The gels were stained with colloidal Coomassie G-250 (PageBlue, Thermo Fischer Scientific) or silver nitrate (57). Protein concentrations were estimated using Coomassie Brilliant Blue G-250 stain calibrated with bovine serum albumin as a standard (58).

